# A phage display library to dissect antibody responses to human coronavirus spike proteins

**DOI:** 10.64898/2026.06.04.730002

**Authors:** Louis J. Taylor, Thiagarajan Venkataraman, Matthew Frieman

## Abstract

Coronaviruses are widespread human pathogens with demonstrated pandemic potential. We developed a phage immunoprecipitation sequencing (PhIP-Seq) library, C-Spike, enabling the profiling of serum antibody responses to coronavirus spike proteins. The C-Spike library includes peptides from 49 *Alpha*- and *Betacoronavirus* spike proteins, including pandemic coronaviruses (SARS-CoV-1, SARS-CoV-2, MERS-CoV), seasonal coronaviruses (HKU1, OC43, 229E, NL63), and selected animal coronaviruses of spillover interest. The library includes a series of 46 amino acid-long peptides covering each spike, with adjacent tiles overlapping by 23 amino acids. Additionally, the library contains alanine-scanned versions of each peptide, tiling three alanine residues at every position across the peptide length, allowing for precise identification of motifs important for antibody binding. We validate C-Spike and its associated bioinformatic analysis pipeline using control antibodies and sera with known reactivity. C-Spike complements existing approaches like neutralization assays to enable deep characterization of antibody responses to coronavirus spike proteins, enabling precise determination of sites important for antibody binding.

## Introduction

Coronaviruses are important pathogens with global impacts on human health. Seasonal coronavirus infections by members of the *Alphacoronavirus* and *Betacoronavirus* genera (229E, NL63, HKU1, OC43) account for an estimated 5-10% of global respiratory disease^1,2^. Additionally, the emergence of highly pathogenic coronaviruses (SARS-CoV-1, MERS-CoV, and SARS-CoV-2) and ongoing discovery of novel animal coronaviruses demonstrate the potential for outbreaks of epidemic and pandemic scales^3–11^. The continued spread and evolution of SARS-CoV-2 to evade pre-existing and vaccine-induced immune responses underscores the importance of understanding antibody responses to coronaviruses and continued development of improved vaccines and therapeutics^12–14^.

Antibody responses to viruses can be tested through multiple laboratory assays involving incubation of isolated antibody or serum with one or more protein substrates, followed by quantification of the resulting interaction. Techniques such as enzyme-linked immunosorbent assays (ELISAs) for specificity, surface plasmon resonance (SPR) for affinity, and neutralization or antibody-dependent cellular cytotoxicity (ADCC) assays for function, yield highly precise information; however, these approaches can be expensive and must be performed individually for different protein targets. A more complete profile of antibody or serum reactivity can be achieved using highly-multiplexed assays such as peptide array or peptide display systems^15,16^.

Phage immunoprecipitation and sequencing (PhIP-seq) is a powerful technique to profile serum antibody repertoires against a variety of proteins simultaneously through creation of libraries of phage displaying peptides derived from proteins of interest^17^. Phage display libraries provide rich data for antibody reactivity across a variety of targets. Additionally, phage libraries are inexpensive; generating more libraries simply requires a bacterial culture. PhIP-seq has been successfully employed to profile antibody responses to pathogens^15,18^, commensal organisms^19,20^, and autoantigens^21^.

Here we introduce C-Spike, a PhIP-seq library for profiling antibody responses to human coronavirus spike proteins. The C-Spike library incorporates peptides from 49 *Alpha*- and *Betacoronavirus* spike proteins, including pandemic coronaviruses (SARS-CoV-1, SARS-CoV-2 and variants, MERS-CoV), seasonal coronaviruses (HKU1, OC43, 229E, NL63), and selected animal coronaviruses of spillover interest. Additionally, C-Spike includes multiple alanine-scanned versions of each peptide with an alanine triplet at every position, enabling precise identification of specific residues important for antibody binding. Spike coverage in the C-Spike library complements existing viral phage display libraries such as VirScan^15,22^ and CoronaScan^23,24^, which contain peptides for other coronavirus proteins (e.g. nucleocapsid and nonstructural proteins), but do not include SARS-CoV-2 variant sequences or full library coverage of alanine-scanned peptides. Overall, the C-Spike library enables precise profiling of antibody repertoires against human coronavirus spike proteins, facilitating improved understanding of humoral responses and enabling development of vaccines with enhanced breadth to control current and future coronavirus outbreaks.

## Methods

### Library design and construction

Forty-nine coronavirus spike sequences were selected from the *Alpha-* and *Betacoronavirus* genera with a focus on known human pathogens, seasonal and epidemic, and animal viruses of zoonotic interest (Table 1). For each spike, tiles of 46 amino acids were generated starting every 23 amino acids (Figure 1), resulting in an overlap of 23 amino acids per peptide with the upstream and downstream peptides. The C-terminal peptide for each sequence was initiated 46 amino acids before the end of the sequence, resulting in an overlap length greater than 23 amino acids between the final and the penultimate peptide. Subsequently, alanine-scanned derivates of each peptide were generated with three alanine residues at every position, resulting in 44 additional alanine-scanned derivatives per library peptide. After adding validated positive and negative control peptides validated to be recognized (or not) by most human sera^22^ and deduplicating peptides with identical sequences, the final library contained 69379 peptides. Reverse translation and generation of adapter-linked oligonucleotides was performed using pepsyn^17^ (https://github.com/lasersonlab/pepsyn). The resulting oligonucleotide library was synthesized by GenScript (Piscataway, NJ, USA).

**Figure 1:**
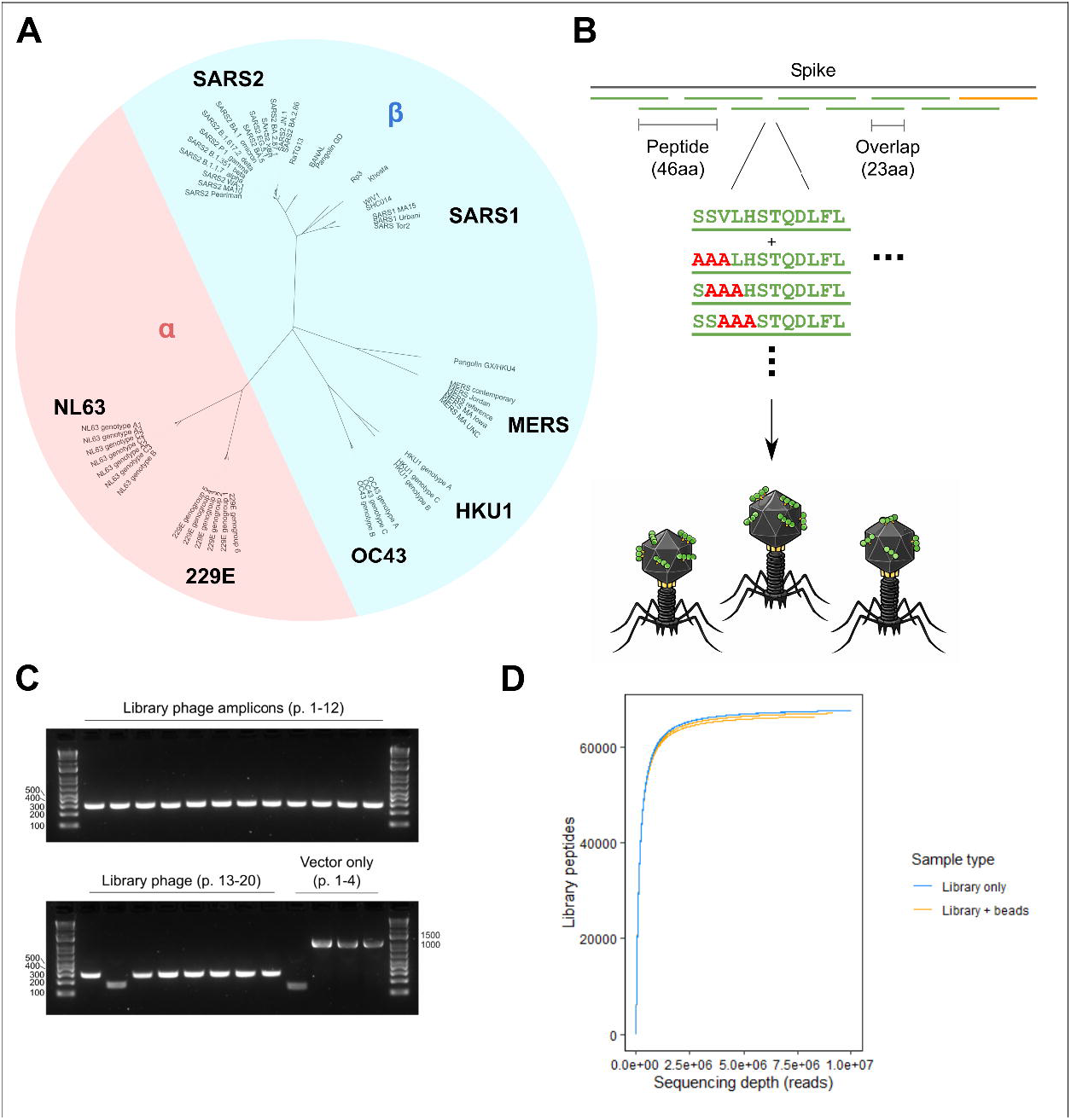
Design of C-Spike library. Design strategy shown in (A). For each spike, peptides of length 46 were tiled every 23 amino acids, resulting in a minimum of 2x coverage of every wild-type library sequence. Additionally, alanine-scanned versions comprising three alanine residues tiled at every position of every wild-type peptide are included in the library. A phylogenetic tree of the 49 spike amino acid sequences included in the library is shown in (B). (C) shows an agarose gel of PCR amplicons from library plaques either from phage vector-plus-library ligation (top gel, left side of bottom gel) or vector-only negative control ligations (right side of bottom gel). Expected product sizes are as follows: successfully cloned library amplicon: 326bp; “empty” (self-ligation): ∼200bp; wild-type vector “stuffer” sequence: ∼1.2kb. (D) shows rarefaction curves for Illumina sequencing of two replicates each of pure library (blue) and library + beads control sample (orange).

The oligonucleotide library was cloned into the T7FNS2 vector^21^ following the protocol detailed in the T7Select® System Manual (MilliporeSigma, Burlington, MA, USA). Briefly, empty vector phage was amplified in *E. coli* strain BLT5403, followed by vector DNA extraction using Qiagen Maxiprep kit (Germantown, MD, USA). Double restriction digest with high-fidelity EcoRI and HindIII was performed, followed by dephosphorylation using quick CIP (New England Biolabs, Ipswich, MA, USA) and gel purification of the vector arms. Restriction sites and adapter sequence were added to the synthesized oligonucleotide library using Herculase II (Agilent, Santa Clara, CA, USA), with EcoRI_PCR1_A5ad_F and HindIII_PCR1_R (primer sequences in Table 2) for 2 cycles, cleaned up using a G50 microspin column (Cytiva, Marlboro, MA, USA), and further amplified with CMV_Fwd and T3 primers for an additional 12 cycles. Products were purified using a Qiagen PCR Cleanup kit (Germantown, MD, USA) followed by double restriction digest with EcoRI and HindIII and gel cleanup. Library-vector ligation was performed at a 3:1 insert-to-vector ratio using T4 DNA ligase (New England Biolabs, Ipswich, MA, USA) then packaged using the T7Select Packaging Kit (MilliporeSigma, Burlington, MA, USA). Seed cultures were amplified in liquid cultured *E. coli* BLT5403 for 4-6h at 37C. Large-scale reagent-grade library was prepared on 150x15mm terrific broth (TB)-agar plates, using a top agar layer of low-melting-point top agarose, 1.6mM Mg^++^, with 1ml of 10x concentrated BLT5403 at optical density (OD) at 600nm of 1.0 for 5-6 h at 37C; phage was harvested when visible clearing of the top agar layer was observed. Phage titering was performed using a similar plate setup, except using 90x15mm LB-agar plates and unconcentrated BLT5403 at OD600 of 1.0. Validation of cloning fidelity and library composition was performed using Sanger sequencing of amplicons generated with T7UP and T7DOWN primers, as well as Illumina sequencing (Figure 1 C-D; Illumina workflow described below).

### Phage immunoprecipitation

Phage immunoprecipitation using the C-SPIKE library was carried out based on the previously published PhIP-seq workflow^17^. Serum volume containing 2 μg of IgG per sample was mixed with 1.48 × 10^10^ total phage in a total of 1ml volume in 96-well deep-well plates, sealed, and rotated for 18h at 4°C. Validation samples tested include anti-S2 commercial antibody NB100-56578 (Novus Biologicals, Centennial, CO, USA); serum from guinea pigs vaccinated with inactivated SARS-CoV (BEI Resources, catalog NR-10361), pooled sera from rabbits inoculated with SARS-CoV spike protein (BEI Resources, catalog NRC-777); human sera from the World Health Organization (WHO) anti SARS-CoV-2 reference panel (NIBSC code 20/268); and in-house naïve mouse serum. Each sample was run in duplicate; sample position on plate was randomized to minimize batch effects. Two wells each were allocated to beads-only and library-only controls. After immunoprecipitation, samples were incubated with 20ul each of protein A and protein G coated Dynabeads (ThermoFisher, Waltham, MA, USA), rotated for 4h at 4°C, then gently washed 3x in 150 mM NaCl, 50 mM Tris-HCL, 0.1% NP-40 (pH 7.5).

Library preparation and barcoding was performed using the standard PhIP protocol^17^ using the primers listed in Table 2. Briefly, for PCR1, thermocycling for 20 cycles (95C initial denaturing for 2min; 20 cycles of PCR [95C denature for 20s, 58C anneal for 30s, 72C extension for 20s]; 72C final extension for 3min), using 0.25uM of PhIP-seq_PCR1_F and PhIP-seq_PCR1_R was performed using Herculase II (Agilent, Santa Clara, CA, USA), followed by barcoding PCR2 using 0.25uM of a unique set of barcoding primers per sample (listed in Table 2). Low primer concentrations were selected to fully deplete primer during thermocycling, enabling pooling of equal volume of PCR product^17^. Pooled PCR products were purified with the QIAquick PCR Purification Kit (Germantown, MD, USA) followed by sequencing on the NovaSeq6000 (Illumina, San Diego, CA, USA) at the Yale Center for Genome Analysis (New Haven, CT, USA).

## Data processing and analysis

PhIP-seq data were analyzed and mapped to peptide sequences and summarized using a Snakemake^25^ pipeline (https://github.com/louiejtaylor/phippy/). The pipeline takes demultiplexed reads as input, performs configurable adapter and quality trimming using cutadapt^26^, counts occurrences of each unique DNA sequence, then maps sequences to either a pre-existing set of peptides or protein database to generate a peptide map *de novo*. Sequence processing uses Biopython v1.86^27^ and pandas v2.3.2^28,29^. The pipeline outputs aggregated counts tables for all peptides in all samples.

Enrichment scores for each sample-peptide combination were calculated by dividing the relative abundance of each peptide in each sample by the mean relative abundance of that peptide in the beads-only control. Data visualization was performed using R v4.4.0^30,31^. Rarefaction curves were generated from peptide read counts from either the library or library plus beads-only samples using the R package vegan v2.6^32^. Spike amino acid sequences were aligned using Clustal Omega v1^33^ and visualized using iTOL v6^34^.

## Results

### Selection of strains for C-Spike library construction

Analysis of coronavirus family members identified forty-nine virus strains in addition to those used in laboratory models (ed. SARS-CoV/MA15^35,36^, MERS-CoV/MA30^37^, SARS-CoV-2/MA10^38^) to be included in the C-Spike library (Figure 1A). The coronaviruses selected are from the Alpha- and Betacoronavirus genera with a focus on known human pathogens and animal viruses of zoonotic interest. Human coronavirus spike proteins selected include seasonal coronaviruses: strains from all 6 genogroups of OC43, 5 genotypes of NL63, 3 genotypes of OC43, and 3 genotypes of HKU1. Strains of animal coronaviruses were selected to include alpha- and betacoronaviruses of relevance to human infection, e.g. published ability to use human angiotensin-converting enzyme 2 (ACE2) to enter cells: RaTG13^39^, BANAL^40^, Rp3^41^, Pangolin GD^42^, Pangolin GX^43^, WIV1^44^, SHC014^45,46^.

For each spike protein, we selected tiled peptides of 46 amino acids long covering each spike from N to C termini, with adjacent tiles overlapping by 23 amino acids (Figure 1B). In addition, alanine-scanned versions comprising three alanine residues tiled at every position of every wild-type peptide are included in the library. Sequences corresponding to peptides were synthesized, subjected to PCR to add restriction cloning adapters, then cloned and packaged into the T7 phage library. Fidelity of packaging was determined by PCR amplification of the inserted spike peptide in phage isolated from plaques from both library and control (vector-only) ligation reactions (Figure 1C). PCR amplification demonstrates correct size of insert in 19 out of 20 plaques from library samples, and Sanger sequencing of these fragments confirmed in-frame insertion of library oligonucleotides.

Following full scale production of the C-Spike library, the library without enrichment was sequenced to confirm equal representation across this region. Two conditions were sequenced in duplicate: library alone, and library subjected to immunoprecipitation conditions without the addition of serum (“library + beads” control)--read counts from the library + beads controls are used as the baseline for calculating peptide enrichment scores. Rarefaction analysis revealed near-complete library coverage at approximately 5 million reads per sample for both conditions (Figure 1D). Together these demonstrate a large and complete library preparation usable for downstream analysis. Once the C-Spike library has been packaged in phage, the library without enrichment was sequenced to confirm equal representation across this region. Sequence analysis shows complete coverage of all spikes in the library with minimal specific peptides overwhelming the library (Figure 1C). Efficiency of packaging was determined by PCR amplification of the inserted Spike gene in isolated phage (Figure 1D). PCR amplification demonstrates correct size of insert in 19 out of 20 isolated plaques. Together this demonstrates a large and complete library preparation for analysis.

### Validation of C-Spike library with known antibodies

The C-Spike library was evaluated with known antibodies to determine functioning of the assay. We first evaluated the performance of the inbuilt positive and negative control peptides using guinea pig, mouse, rabbit and human sera (Figure 2A). The positive control peptide set includes ten peptides from adenovirus, enterovirus, and herpesvirus proteins commonly bound by human serum antibodies, and the negative control set includes ten peptides widely nonreactive across human serum studies^20^. No serum samples enriched for negative control peptides, as predicted. The positive control peptides were widely enriched by human sera but not by rabbit, mouse, or guinea pig sera, consistent with detection of prevalent, public viral epitopes by human serum. These control peptides demonstrate robust, within-sample negative controls for all sample types, as well as an inbuilt positive control for screening human serum samples.

**Figure 2:**
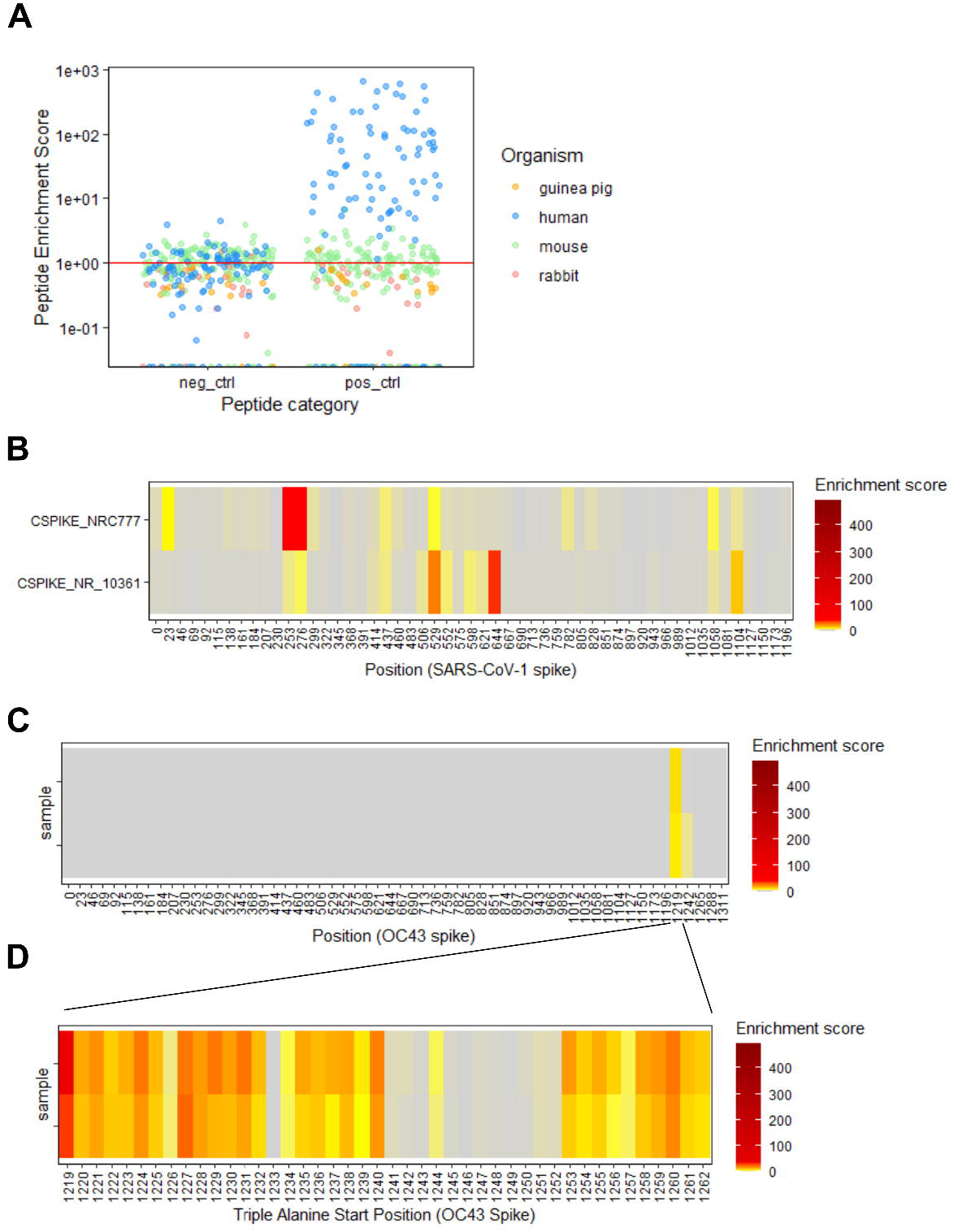
Technical and biological validation. (A) Peptide enrichment scores (relative peptide abundance in experimental samples divided by mean relative peptide abundance in library + beads-only control samples) for library positive and negative control peptides in serum samples from different species. (B) Heatmap showing enrichment scores for SARS-CoV spike reactive guinea pig (NR-10361) and rabbit serum (NRC-777). (C) Heatmap for enrichment scores of a commercial antibody NB100-56578 (Novus Biologicals) raised against a 17-aa peptide towards the 3’ end of the S2 subunit of spike. The two heatmap rows display enrichment scores for both experimental replicates. (D) Heatmap of enrichment scores for alanine-scanned versions of the OC45 peptide beginning at position 1219. The x-axis indicates the position of the first alanine in the triple-alanine series, loss of enrichment at a particular position indicates abrogation of serum antibody binding by alanine mutants. The two heatmap rows display enrichment scores for both experimental replicates.

Next, we evaluated polyclonal sera purified from either rabbits (NR-C777) or guinea pigs (NR-10361) (Figure 2B). NR-10361 is polyclonal guinea pig sera raised against inactivated SARS-CoV-1. Its reactivity is found across multiple epitopes along the Spike protein with hotspots across the Spike protein including the NTD, RBD and S2 region. NR-C777is SARS-CoV-1 Spike polyclonal sera from rabbits. Its reactogenicity is also along the Spike protein with hotspots in the NTD, and S1 region with different epitopes that the NR-C777 sera. This demonstrates that we can profile different reactogenic peptides across diverse sera and Spike proteins depending on viral species, host and platform.

We also evaluated a monoclonal antibody targeting the hCoV-OC43 Spike protein NB100-56578 (Novus Biologicals, Centennial, CO, USA). We find significant reactogenicity against the hCoV-OC43 Spike protein S2 region across duplicate sample preparations (Figure 2C). This demonstrates consistent enrichment of peptides spanning this region and consistency of the assay. We further investigated the precise amino acid binding profile of this monoclonal antibody using the alanine scanning peptides that are a part of the C-Spike library (Figure 2D). The alanine mutations across this region demonstrate where binding is lost for individual peptides that span this region of Spike. We find that the binding is lost (gray bars) when the positions 1241 through 1252 are mutated to alanines. This matches the exact peptide sequence that was used for vaccination of mice to create this antibody. This demonstrates that there is binding of the OC43 monoclonal antibody to this region in Spike.

## Discussion

The C-Spike assay is based on the Phage immunoprecipitation and sequencing (PhIP-Seq) technique which is a high-throughput method to profile antibody binding specificities using phage-display based antigen libraries^17^. PhIP-Seq has been performed using several different antigen libraries (such as viral antigens – all known human viruses, autoantigens – the entire human proteome, environmental antigens – allergens, toxins etc.) to profile humoral responses in different scenarios of health and disease^17,47–49^. The different classes of antigens are designed as peptides of fixed length and cloned into a phage genome to produce T7 phage-display libraries. These libraries are mixed with human or mouse serum and an immunoprecipitation (IP) performed to capture phage/antibody complexes, which are sequenced by next generation sequencing (NGS) to identify antibody specificities.

Our data demonstrate the robust development and practical uses of the C-Spike library for antibody binding site profiling. Using either convalescent sera (Figure 2A) polyclonal sera (Figure 2B) or monoclonal antibodies (Figure 2C), we are able to map the binding site of antibodies across the Spike proteins to specific peptides. Importantly, the alanine scanning peptides that are part of the library, allow for specific binding sites to be identified with single-residue precision (Figure 2D). As each AA is replaced with alanine we can identify which of these changes leads to loss of binding to a specific antibody or serum component.

The limitation of this approach is that all peptides in the library are linear. If an antibody binds to a confirmational epitope that is only present when the full protein is folded correctly, we will not be able to identify their binding site. For those monoclonal antibodies or those present in polyclonal serum, alternative approaches are needed, such as CryoEM mapping. The benefit of PhIP-Seq platform and the C-Spike library is the ability to broadly profile sera across a wide range of Spike proteins with deep coverage and mapping to individual amino acids.

## Figure Legend and Tables

Table 1: Spike amino acid sequences

Table 2: Primer sequences

Table 3: Peptide map

## Acknowledgements

We would like to thank Steve Elledge for providing the BLT5403 phage and phage plasmids for library preparation. Funding for this work is provided by NIH P01AI168347 and NIH 75N93021C00015 / Option 18.1.

